# Multicellular Reservoir Computing

**DOI:** 10.1101/2022.03.26.485905

**Authors:** Vladimir Nikolić, Moriah Echlin, Boris Aguilar, Ilya Shmulevich

**Affiliations:** Bioinformatics Graduate Program, The University of British Columbia, 2329 West Mall, Vancouver, BC Canada V6T 1Z4; Canada’s Michael Smith Genome Sciences Centre at BC Cancer, 100-570 West 7th Avenue, Vancouver, BC Canada V5Z 4S6; Institute for Systems Biology, 401 Terry Ave. N., Seattle, WA 98109, United States; Prostate Cancer Research Center, Faculty of Medicine and Health Technology, Tampere University and Tays Cancer Center, Tampere University Hospital, Tampere, FI-33014, Finland

**Keywords:** reservoir computing, boolean networks, multicellular, simulation

## Abstract

The capacity of cells to process information is currently used to design cell-based tools for ecological, industrial, and biomedical applications such as detecting dangerous chemicals or for bioremediation. In most applications, individual cells are used as the information processing unit. However, single cell engineering is limited by the necessary molecular complexity and the accompanying metabolic burden of synthetic circuits. To overcome these limitations, synthetic biologists have begun engineering multicellular systems that combine cells with designed subfunctions. To further advance information processing in synthetic multicellular systems, we introduce the application of reservoir computing. Reservoir computers (RCs) approximate a temporal signal processing task via a fixed-rule dynamic network (the reservoir) with a regression-based readout. Importantly, RCs eliminate the need of network rewiring, as different tasks can be approximated with the same reservoir. Previous work has already demonstrated the capacity of single cells, as well as populations of neurons, to act as reservoirs. In this work, we extend reservoir computing in multicellular populations with the widespread mechanism of diffusion-based cell-cell signaling. As a proof-of-concept, we simulated a reservoir made of a 3D community of cells communicating via diffusible molecules and used it to approximate two benchmark signal processing tasks, computing median and parity functions from binary input signals. We demonstrate that a diffusion-based multicellular reservoir is a feasible synthetic framework for performing complex temporal computing tasks that provides a computational advantage over single cell reservoirs. We also identified a number of biological properties that can affect the computational performance of these processing systems.

## Introduction

Information processing plays an essential role in cellular systems, enabling cells to adapt their behavior in response to changes in internal and external environmental conditions. By processing information through the heterogeneous, nonlinear interactions of their molecular components, cells can respond to fluctuations in environmental signals (1, 2), search for food (3–5), navigate paths (6, 7), and make behavioral decisions during cellular regulation (8–10). Similarly, when cells aggregate in multicellular communities, the interactions between cells give rise to community-level computation, such as collective decision-making (11, 12), distributed sensing (13, 14), structure formation (15), and immune surveillance (16, 17).

In addition to its importance in natural systems, information processing is playing an increasingly significant role in synthetic biology. In single cell systems, multi-input synthetic molecular circuits are being explored for their potential as sensors, diagnostic tools, and responsive agents for ecological, industrial, and biomedical applications (18). Synthetic intracellular circuits have been implemented in both prokaryotic and eukaryotic cells through a variety of techniques using gene- (19, 20), DNA recombinase- (21, 22), and RNA-based mechanisms (23–25). However, single cell engineering has its limitations (26). Because molecular components cannot be spatially isolated, separate circuits require unique molecular wiring. The number of independent circuits that can reasonably be engineered is therefore limited by the number of manageable molecular species (27, 28). More practically, the complexity of engineered circuits is constrained by the additional metabolic burden they introduce to the cell. Synthetic circuits are also often highly function-specific with limited repurposing (28, 29). To overcome these limitations, synthetic biologists have begun engineering multicellular systems through the combination of simple circuits implemented in individual cells (30). In this way, the burden on individual cells is reduced, subfunctions are separated, and circuit design is modular. Moreover, synthetic biologists can take advantage of other inherent features of multicellular systems, such as the capacity for parallel and distributed computing, robustness to failure, modularity, and scalability (28, 29, 31). Currently, multicellular systems have been designed that compute complex Boolean functions (26, 32–37), act as memory devices (38), behave as glucose sensors (39), generate specified spatial patterns (40), and recognize spatial patterns (41) among other functions (42).

Though different molecular mechanisms are utilized in synthetic multicellular systems, they all rely on intracellular architecture that is designed for a specific function. A flexible computing framework in neural networks, reservoir computing (RC), does not require such specification. Reservoir computing is a signal processing framework in which a signal is input into a fixed-rule dynamic network (the reservoir) and the state of the network is used to continuously approximate a function defined over the input (43, 44). A reservoir computer consists of a signal node(s), a dynamical network (reservoir), and an output node(s). The signal node changes in value over time, passing this value to the reservoir via a subset of reservoir nodes. The reservoir propagates the signal and projects it into a higher dimensional space, retaining a memory of past signal values owing to feedback within the reservoir. Finally, the output node performs a form of regression analysis using another subset of reservoir nodes in order to approximate a given function applied to the input signal over a window of time. Different functions can be approximated simply by retraining the parameters used for regression. Importantly, retraining the output node does not involve any adjustment to the structure or dynamics of the reservoir itself.

Previous work in reservoir computing has shown that a diverse collection of physical systems can be utilized as reservoirs for a range of signal processing tasks, reviewed in (45). In theory, any dynamical system that demonstrates a fading memory property can act as a reservoir (46). Both structurally and dynamically, cells are suitable reservoirs. Intracellular networks are fixed and naturally tuned to balance robustness with responsiveness (i.e. fading memory) (47). Moreover, multiple possible mechanisms - utilizing physical, chemical, and bioelectic mediums - exist for signal input and signal readout. Furthermore, a cellular reservoir computer would be a widely applicable synthetic computing system that would require minimal engineering as the training is performed in the readout layer and not within the reservoir itself. Initial work has already demonstrated the capacity of cells to act as reservoirs. Namely, single cell gene regulatory network reservoirs have been explored in living(48, 49) and theoretical systems (50, 51).

As with other synthetic cellular systems, reservoir computers that take advantage of multicellular populations have the potential for more complex and robust computation compared to single cell reservoirs. Previous work has shown evidence of multicellular reservoirs in the prefrontal cortex (52), cerebellum (53), and glial networks (54). Similarly, in vitro neural systems have been shown to act as reservoirs (55, 56). This work has demonstrated that cells can act together to form a reservoir when connected via neuroelectic signaling. Here, we extend reservoir computing in multicellular populations to the more wide-spread mechanism of diffusion-based signaling, which is used by both prokaryotic and eukaryotic cells and can serve as a means of short- and long-distance communication. Namely, we simulated a proof-of-concept multicellular reservoir computer, in which a 3D community of cells communicating via diffusible molecules serves as the reservoir. We show that multicellular reservoir computing via diffusion-based communication is a feasible synthetic framework for performing complex temporal computing tasks.

## Model description

The goal of the multicellular Reservoir Computer (RC) is to approximate an objective function *f*, given an input sequence of T binary values: *V* = (*v*_1_, *v*_2_,…, *v_T_*), where *v_i_* ∈ {0,1}, 1 ≤ *i* ≤ *T*. The function *f* is defined on a *w* sized window of sequential values from *V* and outputs a binary value: *f*: *X* → *Y*, where *X* = (*v*_*t*–*w*+1_, *v*_*t*-*w*+2_,…, *v_t_*), *w* ≤ *t* ≤ *T* and *Y* ∈ {0,1}. In this work, we have tested the performance of the RC on two time-dependent functions, temporal median and temporal parity, over multiple window sizes. The temporal median and parity functions test the memory and processing power of the reservoir, respectively, and have been used previously for benchmarking reservoir performance (57–60). The median function outputs 1 if there are more 1 bits in the window and 0 otherwise. The parity function outputs 1 if there is an odd number of 1 bits within the window, and 0 otherwise.

### Community organization & communication

The RC is implemented as a community of cells of different strains arranged in a 3-dimensional regular grid of cube shape or in a number of stacked square layers. A visualization of a cell community model can be seen in Figure 1. The cells communicate with each other using Extracellular Signalling Molecules (ESMs). Cells are able to secrete ESMs which diffuse into their surroundings and are able to register the presence of an ESM concentration above a given threshold. The diffusion of ESMs is modeled by solving the standard diffusion equation:

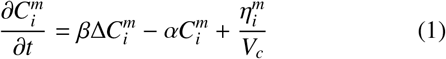

where 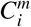 is the concentration of ESM molecule *m* at location *i, β* is the diffusion coefficient, *α* is the degradation rate, 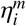 is the secretion rate of molecule *m* by the cell at location *i*, and *V_c_* is the volume occupied by the secreting cell. 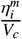 term is 0 if there is no cell at location *i*. *α* and *β* values are constant and same for all ESM molecules. Here we assume that diffusion is a much faster process than regulation of the gene network responsible for secretion and reception, and so we solve the steady state of the equation:

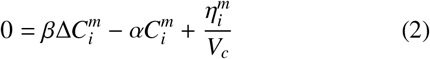

**Figure 1:**
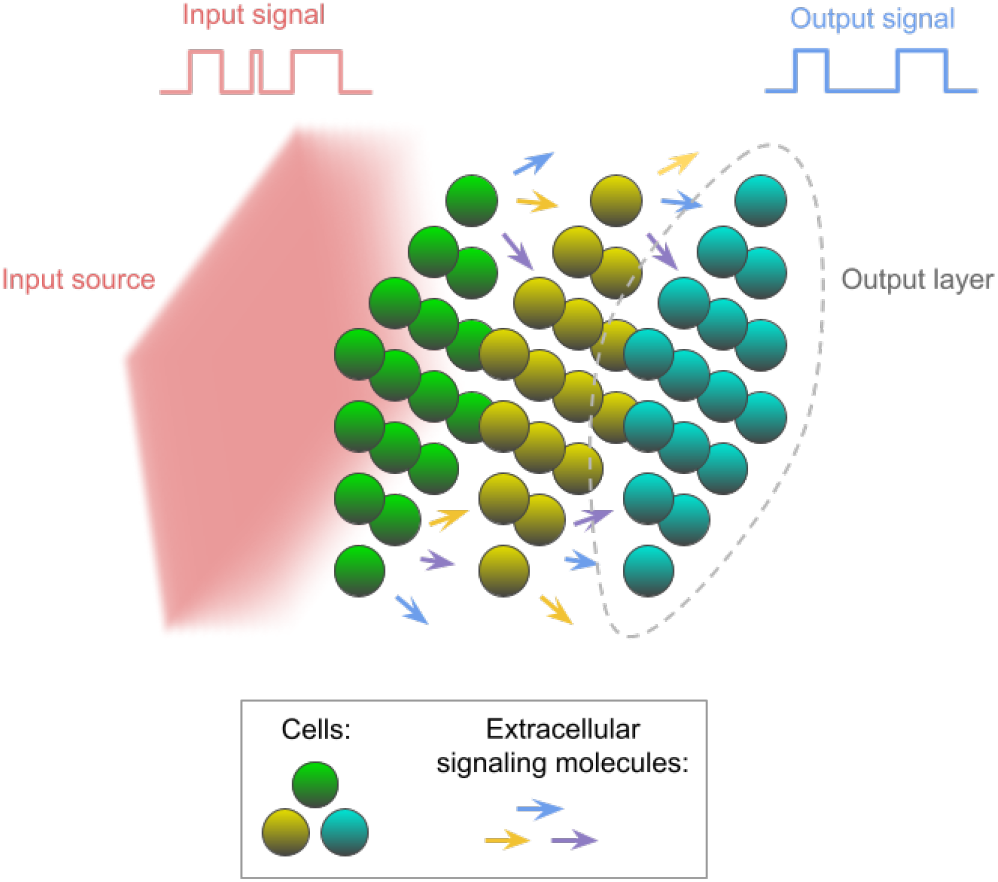
A visual example of a cell community. Cells are arranged in 3D space in a number of square shaped layers along the input signal source axis. The input signal molecules are produced from a simulation domain wall if the value of the input signal in the current simulation time step is 1. The input signal molecules dissipate as they travel through the simulation domain, activating a specified number of cell layers. The cells belonging to these layers register the input and communicate with further layers using ESMs. A specified number of cell community layers at the end opposite of the input source are used for output readout. Their gene values are read at every simulation time step and a regression analysis is performed in order to approximate the objective function.

At the ends of the simulation domain, Neumann boundary condition is enforced with a derivative of 0, i.e. ESMs stay unchanged. This mechanism facilitates information propagation throughout the community.

Each cell has ESM receptor and secreter genes that control communication and can be either ON or OFF. ESM receptors transition into the ON state if the concentration of the corresponding ESM is above the threshold *θ* in the cell’s surroundings and into the OFF state otherwise. The value of *θ* is constant and the same for all ESM molecules. When the secretion genes are in the OFF state, they produce ESMs with a basal secretion rate and diffuse it into the cell’s surroundings. When they are in the ON state, they produce ESMs with an active secretion rate that is higher than basal. For the receptor genes to register ESMs presence, the threshold *θ* is defined as a factor of the basal secretion rate of a single cell. In other words, the ESM concentration should be *θ* times the basal in order to be registered. Incorporating basal secretion in this way allows for a more general model of signaling dynamics better able to capture the leaky nature of real systems (61, 62). Another key parameter of the ESM communication mechanism is the effective interaction distance *λ*, which is derived from the molecule concentration decay rate and the diffusion coefficient: 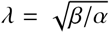. Increasing λ allows ESMs to travel further from their source, enabling more distant cells to communicate with each other.

### Cell inner workings

Inside each cell, gene regulation is modeled with a Random Boolean Network (RBN) (63) implemented as a graph *G* = {*G_V_*, *G_E_*}, where *G_V_* is a set of *N* nodes/genes: *G_V_* = {*g*_1_, *g*_2_, …, *g_N_*}, and *G_E_* is a set of 2 · (*N* – 1 – *E*) edges, where *E* is the number of ESMs. Each gene can either be OFF or ON: *g_i_* ∈ {0,1}, 1 ≤ *i* ≤ *N*. All cells have the same number of genes defined as a simulation parameter (*N*). There are four types of genes in each cell: input signal gene, ESM receptors, ESM secreters, and reservoir genes. There is always a single input signal gene (1), a receptor and a secreter gene for each ESM (2 · *E*), and the rest are reservoir genes (*N_r_*), for a total of *N* genes. While the input and ESM genes are responsible for external communication, the reservoir genes represent the internal state of a cell.

A simulation is divided into discrete time steps. At every time step *t*, the input signal is diffused into the environment (as described in the following subsection) and the ESMs secreted based on secreter gene states. Then Equation (2) is solved for every location in the simulation to determine molecule concentrations. Following this, the input signal gene *g*_1_ and the ESM receptor genes determine their states based on the presence of the input signal and ESM molecules respectively in the cell’s surroundings. For a corresponding gene *g_i_,* if the molecule concentration is above a threshold, then *g_i_*(*t*) = 1 and *g_i_*(*t*) = 0 otherwise. The input signal gene *g*_1_ has a different threshold from ESM receptors, as described in the following subsection. Finally, ESM secretion genes and reservoir genes determine their values based on the state of the other genes. The value of each of those genes, *g_i_*(*t*), is updated using a function *f_i_*. *f_i_* has *k_i_* inputs chosen randomly and uniformly from *G* excluding ESM secreter genes, *g_i_*(*t* + 1) = *f_i_*(*g*_*j*1_(*t*), *g*_*j*2_(*t*),…, *g_ki_*(*t*)), *g_j_* ∈ *G* \ *G_ESMsecreters_*. The fraction of genes whose *f_i_* inputs include the input signal gene *g*_1_ is controlled with the *L* simulation parameter. The truth table for *f_i_*, its output for each possible combination of input values, is randomly generated. The average number of inputs (in-degree) for all *f_i_* is 2 and the probability of *f_i_* outputting a 1 is 0.5. These function parameters have been shown to maximize computing capacity (64, 65).

In each simulation, not all cells are described by the same set of functions. Each cell is randomly assigned a strain. Each strain has a distinct set of randomly generated edges in *G*, defining its RBN topology. and a distinct truth table is generated for each *gi.* The number of strains is a simulation parameter (*S*).

### Community input/output

A random input signal sequence *V* is generated for each simulation. For every time step *t*, there is a corresponding input value *v_t_*. The input signal is represented as molecules diffusing away from one of the simulation walls, as shown on the left part of Figure 1. The dynamics of the input signal molecules are modeled using the same diffusion equation as the ESMs. The difference is that the secretion term is always 0 and instead the production is modeled with Dirichlet boundary condition on the signal producing wall. All other walls still follow Neumann boundary condition with a derivative of 0. If the input value *v_i_* is 1 in a simulation time step, the solution to the Dirichlet boundary condition is >0 (100,000), diffusing the molecules from the simulation wall into the environment, affecting nearby cells. The input molecules move towards the opposite wall of the simulation domain, dissipating along the way and penetrating *I* layers of the community, specified as a parameter. Each simulation calculates what threshold cells should have to the input signal in order for it to penetrate *I* layers. If *v_i_* is 0 in a simulation time step, the Dirichlet boundary condition solution is 0 and no input signal molecule is produced.

As information is processed by the community, the goal is to obtain an approximation of the given function *f* (median or parity) on the input signal by reading the cell states of the cell community output layers. A number of cell community layers, specified as a parameter (*O*), on the side opposite of the input signal producing wall are used for output. After the ground truth for *f* output is calculated, the gene states *g_i_* of all cells within the output layers are used as covariates for regression analysis with the ground truth values as desired output. Further description of the reservoir training and testing are provided in Materials and Methods.

## Results

Here we have tested the performance of a cube-structured community as a reservoir with comparison to a single cell (Proof of Concept); analyzed the effect of parameter values on the cube-structured reservoir’s performance (Sensitivity Analysis); and tested the performance of a 3-layered rather than cube-structured community of cells as a reservoir (Layered Community). In all analyses, accuracy has been used as a metric of performance. We define accuracy as the ratio of correctly approximated outputs using the regression model over the total number of outputs. Since the tested functions have binary output, at worst, the performance is 50%, which is equal to a random guess, and 100% at best, where all outputs are correctly guessed.

### Proof of Concept

In this analysis, we simulated a cube-shaped community in which all cells are exposed to the input signal and all cells are used as outputs. In this way, we focused on the multicellular and communication aspects of the population rather than the effects of spatial heterogeneity in the input signal (input heterogeneity). The cube-shaped community was tested for approximating median and parity functions with window sizes 3, 5, 7, and 9. We also simulated a single cell to compare multicellular to unicellular performance. The fixed simulation parameters can be found in Supplementary Table S1, with the only difference between the multicellular and the unicellular system being the number of cells, 12^3^ and 1 respectively. The performance comparison can be seen in Figure 2.

**Figure 2:**
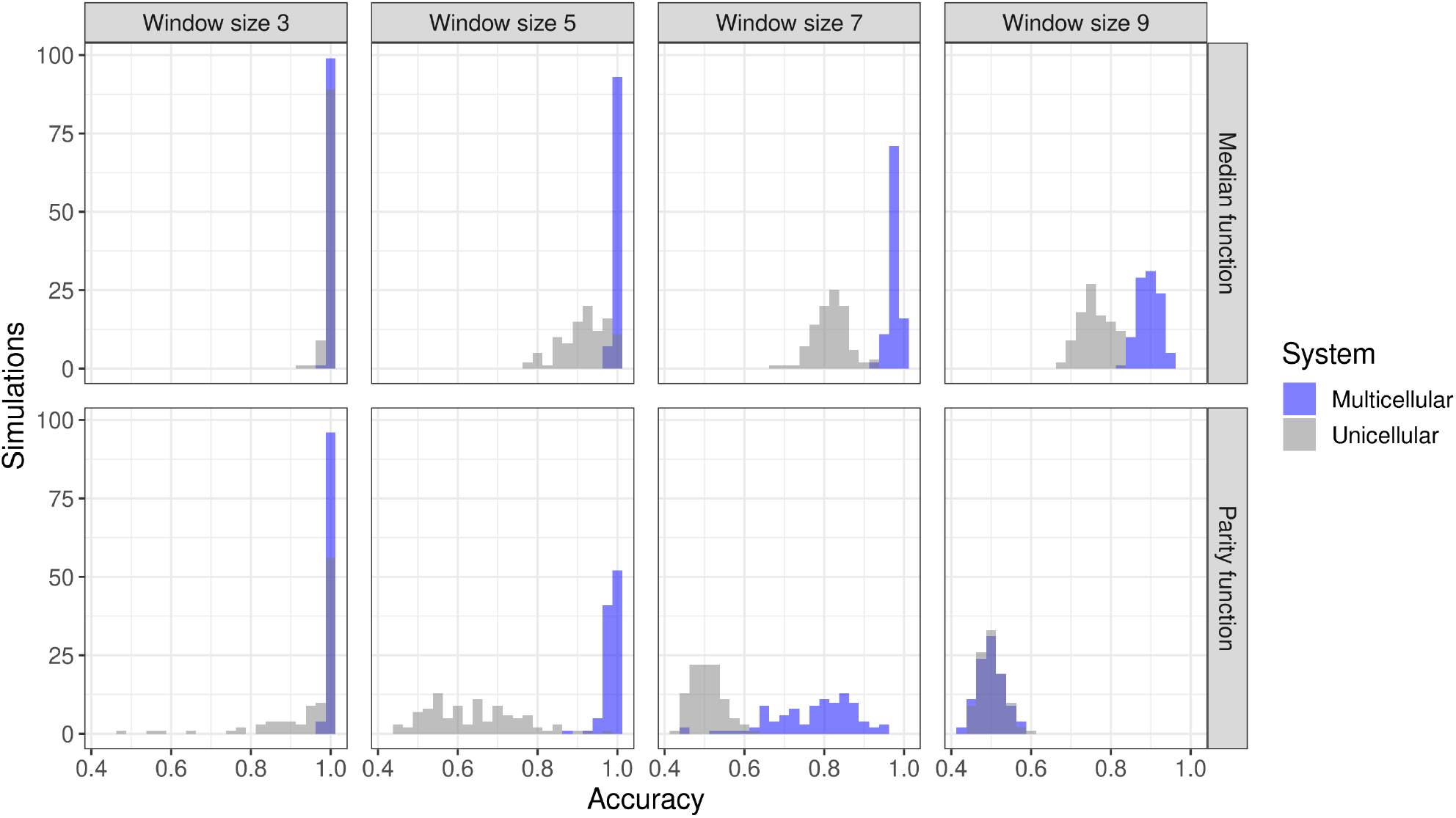
Simulation results from the Proof of Concept analysis. The distribution of reservoir accuracy is shown for median and parity functions with different function window sizes. To generate the distributions, each combination of window size and function was simulated 100 times. As parity is a harder function to approximate, the distributions are skewed towards the left compared to median function. For window size 3, both functions are well approximated, with almost all simulations performing with 100% accuracy. As window size increases, distributions move towards 50% (random chance) accuracy, with parity function reaching around 50% for window size 9 for both unicellular and multicellular systems. These results demonstrate that a community of cells can act as an RC and can outperform a single cell in that capacity.

The results demonstrate that a community of cells has the capacity to approximate binary functions even with larger window sizes. For estimating the median function, the mean accuracy remains near 1 for all window sizes tested, but does decrease as the window size increases. For estimating the parity function, a computationally harder task, the community performs nearly perfectly for window sizes 3 and 5, but mean accuracy rapidly decreases for larger window sizes. At window size 9, mean accuracy is around 50%, or as good as a random guess, demonstrating the computational capacity limit of this setup. In all cases except parity window size 9 where both systems are at their limit, multicellular communities achieve a higher average accuracy when compared to the performance of a comparable unicellular reservoir simulated here as well as to previous reports of single cell Boolean Network (BN) reservoirs (50).

### Sensitivity Analysis

To examine how community parameters, especially those related to intercellular communication, affect reservoir performance, sensitivity analysis was performed using Partial Rank Correlation Coefficient (PRCC) (66). Briefly, sensitivity analysis generates random sets of input parameters and assesses the accuracy of the model for every parameter set in order to determine the rank correlation between each parameter *x_i_*, 1 ≤ *i* ≤ *k*, where *k* is the number of parameters, and the output *y* (Accuracy). Correlation is calculated after rank-transforming values, and so non-linear effects are ignored. When calculating correlation for each parameter, the effect of other parameters is cancelled out (further explained in Materials and Methods).

The model was tested to approximate parity function with a window size of 7. The community is cube-shaped and all cells are used for output. The results of Sensitivity Analysis can be found in Table 1, including the range within which parameters have been randomly sampled. The rest of the parameters are kept fixed and can be found in Supplementary Table S2.

**Table 1:** Sensitivity Analysis results, including parameter range, rank correlation (*ρ_R_*), and *p-value.* As shown in earlier works, *L* and *N_r_* have strong correlation with reservoir accuracy. These results add *C* and *S* to the set of parameters that are strongly correlated with accuracy. The number of input layers, *I,* the number of ESMs, *λ*, and *θ* all have noticeably lower correlation.

| Parameter | Range | $\rho_R$ | $p$-value |
| --- | --- | --- | --- |
| Gene input fraction ($L$) | [0.1, 1.0] | 0.77 | $< 1.00e - 10$ |
| Reservoir genes ($N_r$) | [10, 500] | 0.55 | $< 1.00e - 10$ |
| Cells ($C$) | [1, $15^3$] | 0.42 | $< 1.00e - 10$ |
| Strains ($S$) | [1, 20] | 0.33 | $< 1.00e - 10$ |
| Input layers ($I$) | [1, 15] | 0.08 | $< 1.00e - 10$ |
| ESM $\lambda$ | [2.5, 30] | 0.04 | $1.63e - 05$ |
| ESMs ($E$) | [0, 20] | -0.06 | $4.54e - 09$ |
| ESM $\theta$ | [10.0, 100.0] | -0.06 | $1.08e - 09$ |

As demonstrated in previous RBN reservoir studies (50, 57), the fraction of genes connected to the input gene (*L*) and the number of reservoir genes (*N_r_*) strongly positively correlate with accuracy (*ρ*_*R*(*L*),*R*(*y*)_ = 0.77, *ρ*_*R*(*N_r_*),*R*(*y*)_ = 0.55). Here, we demonstrate that the number of cells (*C*) and the number of strains (*S*) also positively correlate with accuracy (*ρ*_*R*(*C*),*R*(*y*)_ = 0.42, *ρ*_*R*(*S*),*R*(*y*)_ = 0.33). Additionally, the number of cell layers that receive the input signal (*I*) moderately positively correlates with accuracy (*ρ*_*R*(*I*),*R*(*y*)_ = 0.08). Despite being responsible for intercellular communication, the number of ESMs (*E*), *λ*, and *θ* only have a weak correlation (*ρ*_*R*(*E*),*R*(*y*)_ = 0.04, *ρ*_*R*(*θ*),*R*(*y*)_ = −0·06).

### Layered Community

This analysis involves a cell community of three square shaped layers along the input signal axis, as depicted in Figure 1. The model was tested to approximate parity function with a window size of 3. This function and window size were chosen because the difficulty of the task balances the accuracy well between 50% and 100%. In this setup, the threshold for the input signal is such that the signal is undetectable past the first layer. Additionally, only the last cell layer is used for output. The ESM *λ* parameter value is such that the signal is only detectable to immediately neighbouring cells in the same or adjacent layer. Because of this, the first layer cannot directly communicate with the third output layer. The information has to propagate through the middle layer, and so it takes two simulation time steps to reach the output layer. The intention here is to model the input heterogeneity present in real cellular communities, which can be the result of receptor heterogeneity, limited signal penetration, or engineered spatial segregation. Moreover, With this setup, we more closely tested the role of communication in the reservoir, since the output layer relies solely on intercellular communication for information about the input signal. An analysis of the average performance of the Layered Community for median and parity functions shows that reservoir performance is generally improved over the performance of single cells (Supplementary Figure S4).

Four parameters have been varied in order to determine their effect: number of reservoir genes (*N_r_*), cells (*C*), strains (*S*), ESMs (*E*). *N_r_* is relevant to the relationship between individual computational power and community computational power; C and S are relevant to the size and diversity of the reservoir; and *E* is relevant to the informational bandwidth of communication. The numbers for cells were chosen such that they can form three square layers without excess cells. Figure 3 shows the relationship between each parameter and accuracy, while keeping all other parameter values fixed (Supplementary Table S3). Each datapoint is the average accuracy of 100 simulations.

**Figure 3:**
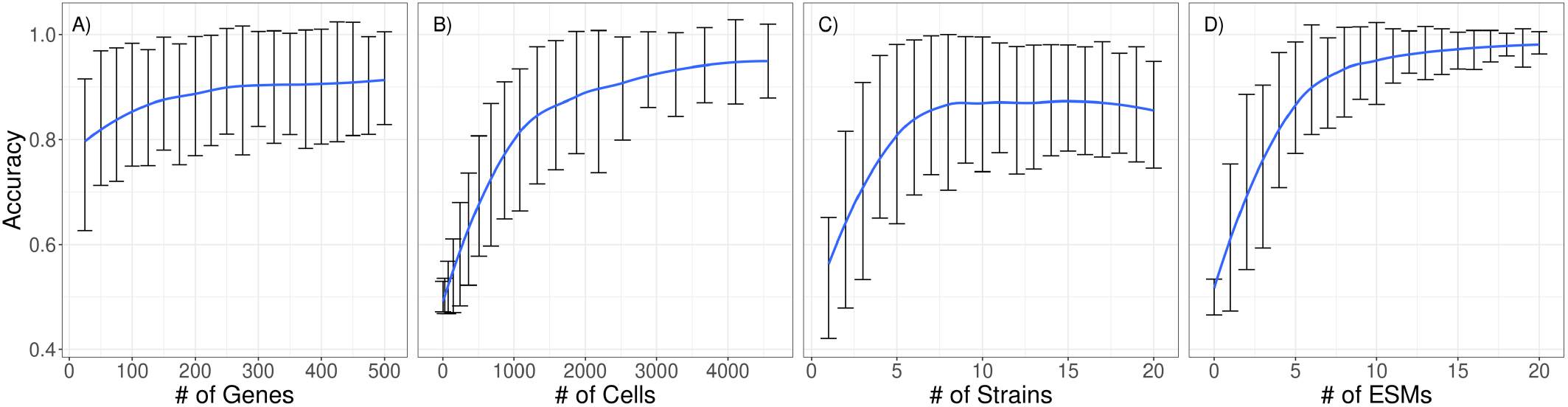
Simulation results from the Layered Community analysis. Four parameters were examined independently while keeping the rest fixed: the number of genes (A), cells (B), strains (C), and ESMs (D). The blue line shows the trend in average accuracy. Each data point along the trend line is the average accuracy of 100 simulations and the bars show ± one standard deviation. This data shows that the number of cells and ESMs contribute the most to improving accuracy. These are also the parameters most closely responsible for carrying the signal across layers. Genes give a modest improvement to accuracy and plateau around 300. Finally, strains initially improve accuracy noticeably, but peak around 8, after which accuracy starts decreasing.

With the exception of the number of strains, all parameters have a monotonic positive relationship with accuracy. For the number of genes and cells, this aligns with the conclusion from Sensitivity Analysis, i.e., that they positively correlate with accuracy. However, gene contribution to accuracy plateaus after around 300 genes. Strains have also been found in Sensitivity Analysis to correlate positively, but in this setup plateau quickly and do not have a clearly monotonic relationship as they start to negatively correlate with accuracy after around 8 strains. Another contrast from Sensitivity Analysis is the clear contribution of the number of ESMs to accuracy. Unlike in Sensitivity Analysis, here the output cells did not have direct access to the input signal and relied on ESM communication through layers.

## Discussion

With this work, we aimed to demonstrate that cellular communities utilizing diffusion-based communication are capable of acting as reservoir computers. Previous research has successfully applied neuronal communities with neuroelectric communication schemes as reservoirs. However, communities that rely on diffusion, a more widespread form of communication, had yet to be tested in this role. Here we have provided theoretical evidence demonstrating the potential of such reservoirs. We further demonstrated that these reservoirs are successful under conditions of both input homogeneity, when all cells have access to the input signal and can act as individual reservoirs, and input heterogeneity, when only some cells have access to the input signal and communication is necessary to function as a reservoir. The latter case is likely to be more applicable to engineered systems in which input heterogeneity can be introduced through receptor expression heterogeneity, reduced signal penetration, or externally imposed spatial segregation.

In addition to demonstrating that multicellular communities can act as reservoirs, we have shown that they can significantly outperform their single cell counterparts. Not only was average accuracy higher, but cellular communities were able to perform tasks that single cells could not (Figure 2, parity function window size 5 and 7). However, the factors responsible for the increased computational power remain unclear. Two potential factors are the increased number of individually capable reservoir computers (cells), from which a prediction can be crowd-sourced; and the increased size of the total cellular network through communication, which has more components to contribute to the computing capacity. We found that removing communication when it is not necessary has little effect on performance (Supplementary Figure S2). Thus, crowd-sourcing from many cells and strains of cells likely plays a significant role in the success of multicellular reservoirs. Our brief analysis also suggests that communication can negatively impact accuracy when little memory is required for the signal processing task and positively impact accuracy when more memory is required (Supplementary Figure S3, median function). Therefore, communication may be increasing the memory of the system as a whole while also reducing the independence of each cell, thereby reducing the value of crowd-sourcing.

For reservoirs that rely on communication to propagate the input signal, we found that performance can also exceed that of single cells (Supplementary Figures S3 and S4); though we did observe a decrease in reservoir performance compared to reservoirs that do not need communication. Estimation of the parity function is also notably more difficult when communication is required. The inherent delay as the signal is propagated from input to output cells may be causing the decrease in performance. Previous reservoir computing research has shown that introducing a delay between receiving an input and estimating the function negatively affects performance (57). Another possibility is that signal propagation between cells introduces noise and uncertainty to the signal, which would contribute to lower accuracy. Unexpectedly, the spatial organization of input and output cells has a very strong effect on reservoir performance. Populations with one half of cells receiving input and the other half used as output, randomly intermixed in the community, are at a disadvantage (Supplementary Figure S3). Intermixed populations actually increase in accuracy as window size increases. More targeted experimentation is needed to further explore the role of communication in reservoir behavior.

A multicellular reservoir’s computing capacity also varies with properties inherent to both the individual cell and the cellular community. Here we focused on parameters that have the potential to be incorporated into the design of a multicellular reservoir computer. As previous work has shown for single cell RBN reservoirs, the number of genes within each cell and the number of those genes with input from the signal both positively correlate with the performance of multicellular reservoirs (50, 57). Intuitively, more genes means more computing power and more genes wired to the input signal means less of an information bottleneck. Notably, there is a nontrivial amount of heterogeneity across reservoir performance with respect to the number of genes and the benefit of adding more genes to each cell quickly diminishes (Figure 3). For engineered systems this suggests that the burden to an individual cell can be kept low without losing out on much computational capacity.

Considering properties unique to cellular communities, we found that the size, diversity, and number of signaling channels of the community positively correlate with reservoir performance. If crowd-sourcing is, in fact, a contributing factor to computing, then increasing the number and diversity of cells would be advantageous. The effect of ESMs is less straightforward. Accuracy in the communities that do not require communication (Proof of Concept) shows little correlation with ESM-related parameters, suggestion a minor improvement that communication provides to these reservoirs. Although, it is also possible that the method used for sensitivity analysis is simply unable to detect a relationships since the interaction distance and threshold form a nonlinear communication parameter space. For the three-layered community, in which communication is necessary, increasing the number of signaling molecules has a large effect on accuracy, though the reason is not obvious. While more signaling molecules may benefit the efficacy of signal transmission, it would also reduce the independence of individual cells, which may negatively impact the reservoir. Overall, it appears that a minimum number of cells, strains, and ESMs is necessary for reservoirs to perform well, but that deficiency in one parameter can likely be compensated for by increasing another.

Though we have tried to consider real-world conditions in this work, further work is required to identify all limitations that could be encountered in the bio-engineering process. For example, the number of genes that can directly interact with signal receptors may be severely limited in practice. Additional analysis to account for such limitations would be straightforward to test in a similar computational framework. Future work could also include a closer investigation of the effects of communication on computational power versus system memory as well as the effect of different communication regions described by the communication parameters (Supplementary Figure S7).

What we have shown with this work is that multicellular communities can act as reservoirs. We have identified that the communication scheme and community structure both play important roles in the type of computation performed. And lastly we have shown that, in addition to the benefits typically associated with multicellular systems, cellular communities can easily outperform single cells with more flexibility in how they can be optimized.

## Methods and Materials

The simulations were implemented using the Biocellion v1.2 multicellular simulation framework (67). Fixed simulation parameters can be found in Supplementary Tables S1 to S3 and their meaning is further described here.

### Simulation details

All simulations run for *T* = 1000 time steps. Before each simulation, a random binary input signal array *V* is generated, equal in size to the number of simulation time steps. In each time step *t*, the input signal molecule is produced if *v_t_* is 1 (Dirichlet boundary condition is 100,000), otherwise it is not (Dirichlet boundary condition is 0). A numerical solution is implemented that determines the threshold each cell should have to the input signal when it is being produced so that number of cell layers that detect the presence of the input signal molecule is equal to the value of the input layers *I* parameter. All cells that belong to the input layers register the input signal presence and set the state of the input gene *g*_1_ to 1 for the duration of that simulation time step. The state of *g*_1_ is 0 otherwise. For all simulations, *v*_0_ is guaranteed to be 0 and all molecule concentrations are initialized to 0. The input signal gene *g*_1_ and ESM receptor genes are initialized to 0 for all cells. The initial values of the rest of the genes are randomly generated.

For Proof of Concept and Sensitivity Analysis, cells are arranged in a cube shaped grid, whereas for the Layered Community setup, the cells are arranged in a grid of three square-shaped layers. Each cell occupies one simulation domain voxel and is centered in it.

### Random boolean networks

Every cell has *N* genes: one input signal gene (1), one receptor and one secreter gene for each ESM (2 · *E*) and the rest are reservoir genes (*N_r_*). When assigning edges in the RBN, the constraints are:

- Input gene and ESM receptor genes cannot have incoming edges.
- ESM secreter genes cannot have outgoing edges.

In order to stay within these constraints and have an average in-degree of two for the reservoir genes, there is a total of 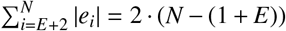 randomly and uniformly assigned edges in the RBN, where |*e_i_*| is the in-degree of gene *i*. The incoming edges are counted from *E*+2 because the input gene (1) and the ESM receptor genes (*E*) do not have incoming edges. A fraction of these edges is used specifically for connections with the input gene. At most, there are *E* + *N_r_* = *N* – (1 + *E*) edges from the input gene (as the other 1 + *E* genes do not have incoming edges). The number of these edges is *L* · (*E* + *N_r_*), where 0 ≤ *L* ≤ 1 is a parameter. Additional information about the RBN construction within cells can be found in Supplementary Random boolean network construction section.

Cells communicate with each other using ESMs, which are implemented using the diffusion mechanism in Biocellion. Here we assume that molecule diffusion is a much faster process than gene regulation and so diffusion is implemented with a steady state partial differential equation (PDE). A single simulation time step is sufficient for both the input signal and ESMs to reach the steady state. Following this, the RBN for each cell is updated. For each ESM molecule *m,* diffusion in the environment follows Equation (2) where 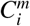 is the concentration of ESM molecule *m* at voxel *i, β* is the diffusion coefficient, *α* is the degradation rate, 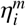 is the secretion rate of molecule *m* by the cell at voxel *i*, and *V_c_* is voxel volume (all voxels have the same volume).

This PDE is numerically calculated (68, 69). We also define *λ* as the effective interaction distance, where 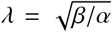. The *θ* parameter is the threshold for registering ESM presence in the cell’s voxel. It is defined as a factor of the ESM concentration in a cell’s voxel when the cell is producing the ESM molecule at basal rate and this concentration is numerically calculated. The same value of *θ* is used for all the ESM. For Proof of Concept and Layered Community, the ESM communication parameters: *α* (1.0/45.0), *β* (5.0), λ (15.0, derived from *α* and *β*), *η* (1.0 basal, 5.0 active), and *θ* (11.5) have the same values as in the previous work done by Echlin et al (50). For Sensitivity Analysis, *β* and η are the same as for Proof of Concept and Layered Community, whereas λ and *θ* parameter ranges are informed by Supplementary Figure S7. Based on information from previous work (70), the tested parameter values explore scenarios where:

- Intercellular communication does not occur, because the receptor is always on, always off, or only responds to self-generated signal.
- Intercellular communication occurs, where the receptors either respond to neighbor-generated signal, self-generated signal, or a combination of the two.

For each simulation time step, actions are done in the following order (as implemented in Biocellion):

1. ESMs are secreted into the environment based on secreter gene states.
2. Input signal Dirichlet boundary value is set to 100,000 if the current input signal value is 1, and to 0 otherwise.
3. PDEs are solved and molecule concentrations in every voxel stabilized.
4. Gene values for all cells are updated. Input gene and receptors according to the environment, and secreters and reservoir genes according to the update functions.

### Linear Regression

Linear regression analysis was done using the LassoCV function (with *selection* = “*random*”, *tol* = 0.05 parameters for faster convergence) from the Python language scikit-learn library (71). Each simulation starts off with *T_W_* = 100 warm-up time steps, included in the total *T* = 1000 time steps, during which the simulation runs as normal, but the outputs are not used for linear regression. The covariates of linear regression are all reservoir gene states of all cells belonging to the output layers, which are collectively denoted here as *g^O^*. The number of output layers (*O*) is a parameter and the layers are selected sequentially from the community, starting at the side opposite of the input signal source. Additionally, we define the parameter *d* which is the delay in time steps between the objective function ground truth output at time *t*, *y_t_*, and *g^O^* which is taken from time *t* + *d*, denoted as *g^O^*(*t* + *d*). This delay parameter lets the information propagate through the system for *d* timesteps before linear regression is performed. This is necessary in Layered Community, where it takes two time steps for the signal to propagate from the first to the last, third community layer. For that reason, *d* is 2 in that analysis. The input to linear regression is matrix *X*, where columns correspond to covariates/genes and rows to time steps. If *X_i_* is the *i^th^* row of *X* and *w* the objective function window size, then for each time step *t, T_W_* + *w* + *d* ≤ *t* ≤ *T*, *X*_*t*–*T_w_*+*w*+*d*)+1_ = *g^O^*(*t*). Additionally, the ground truth array *Y* for the objective function *f* is calculated for each time step *t*, *T_w_* + *w* ≤ *t* ≤ *T* – *d*, on the input sequence window: *Y*_*t*–(*T_w_*+*w*)+1_ = *f*(*V*_*t*–*w*+1_, *V*_*t*–*w*+2_,…, *V_t_*). The rows of *X* and *Y* are divided into training (75%) and test dataset (25%) randomly. LassoCV is ran on the training dataset and the accuracy of the model is assessed on the test dataset.

### Sensitivity analysis

For Sensitivity Analysis, Latin Hypercube Sampling (LHS) (72) is used to generate 10,000 samples, each simulated 5 times. For each sampled parameter, LHS divides the sampled range into 10,000 equal intervals and takes a randomly selected value from each interval. This ensures a fairly uniform sampling which would not be guaranteed by brute force sampling. The order of selected values is then randomly shuffled for each parameter.

In order to calculate rank correlation in Sensitivity Analysis for each parameter individually, the effect of other parameters has to be cancelled out. The parameters and output are first rank-transformed to ignore non-linear effects and keep only monotonicity. Then, for every rank transformed parameter 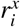 and output *r^y^*, the following linear models are built that estimate the parameter and output from other parameters:

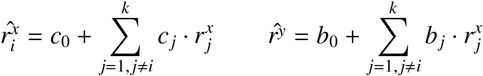

Pearson correlation is then calculated between the residuals, which have linear effects of other parameters removed: 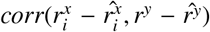. The plots showing correlation between each varied parameter and accuracy can be seen in Supplementary Figure S5, and partial correlation after rank transformation in Supplementary Figure S6. The distribution of accuracies for Sensitivity Analysis simulation runs can be seen in Supplementary Figure S1. The fixed parameters for Sensitivity Analysis, found in Supplementary Table S2 were chosen such that the accuracy distribution would be evenly spread out. If too many simulations were concentrated at 0.5 or 1.0 accuracies, the Sensitivity Analysis results would be less reliable.

### Miscellaneous

Plots in this work were generated using the ggplot2 (73) and seaborn (74) libraries.

The code used in this work can be found online at: https://github.com/vlad0x00/multicell-rc

## Acknowledgments

V.N. was supported in part by a scholarship from the UBC Bioinformatics Graduate Program via the NSERC CREATE program in High-Dimensional Biology. The authors wish to acknowledge Canada’s Michael Smith Genome Sciences Centre, Vancouver, Canada for computing services. The GSC’s IT infrastructure is supported by WestGrid (www.westgrid.ca) and Compute Canada (www.computecanada.ca). M.E. acknowledges support from Tampere University, Tampere, Finland. I.S. and B.A. gratefully acknowledge support from the ISB.

## Author contributions

Project design (VN, ME, BA, IS), Simulation (VN), Manuscript preparation (VN, ME, BA, IS)

