## Supplementary Data for "Multicellular Reservoir Computing"

#### Random boolean network construction

The total number of edges in a Random Boolean Network (RBN) within a cell is:

$$e = (N - (1 + E)) \cdot 2$$

where $N$ is the total number of genes and $E$ the number of Extracellular Signalling Molecules (ESMs), which is equal to the number of ESM receptor genes. The $1 + E$ part comes from the fact that the input gene (1) and ESM receptors ($E$) do not have incoming edges. The rest of the genes should have an average in-degree of 2, hence the factor at the end. From $e$, a subset of edges are used for connections with the input gene. The number of these input edges is modulated with the $L$ parameter, $0 \leq L \leq 1$. The number of input edges is:

$$e_i = (N - (1 + E)) \cdot L$$

To assign the edges to nodes, first, two sets of edges are generated:

1. All possible edges that start at the input gene.
2. All possible edges that do not start at the input gene.

The generated edges exclude those that end in the input gene or ESM receptors. These two sets are randomly shuffled, and from the first set, the first $e_i = (N - (1 + E)) \cdot L$ are selected and assigned. From the second set, the first $e - e_i = (N - (1 + E)) \cdot 2 - (N - (1 + E)) \cdot L$ are selected and assigned. This way, we randomly and uniformly assign edges with an average in-degree of 2 and the exact number of input edges we wanted.

#### Supplementary tables

Table S1: Proof of Concept fixed parameters. Cells are arranged in a $12x12x12$ cube. All cells are reached by the input signal and all cells are used for output.

| **Parameter** | **Value** |
|---|---|
| Gene input fraction ($L$) | 0.5 |
| Reservoir genes ($N_r$) | 100 |
| Cells ($C$) | $1728(12 \cdot 12 \cdot 12)$ |
| Strains ($S$) | 10 |
| Input layers ($I$) | *All* |
| Output layers ($O$) | *All* |
| ESM interaction distance ($\lambda$) | 15.0 |
| ESMs ($E$) | 5 |
| ESM threshold ($\theta$) | 11.5 |
| Delay ($d$) | 0 |
| ESM diffusion coefficient ($\beta$) | 5.0 |
| ESM basal secretion rate | 1.0 |
| ESM active secretion rate | 5.0 |
| Cell radius | 10.0 |
| Voxel side length | 20.0 |
| Warm-up time steps ($T_W$) | 100 |
| Total time steps ($T$) | 1000 |

Table S2: Sensitivity Analysis fixed parameters. All cells are used for output.

| **Parameter** | **Value** |
|---|---|
| Output layers ($O$) | *All* |
| Delay ($d$) | 0 |
| ESM diffusion coefficient ($\beta$) | 5.0 |
| ESM basal secretion rate | 1.0 |
| ESM active secretion rate | 5.0 |
| Cell radius | 10.0 |
| Voxel side length | 20.0 |
| Warm-up time steps ($T_W$) | 100 |
| Total time steps ($T$) | 1000 |
| Function | *parity* |
| Window size ($w$) | 7 |

Table S3: Layered Community fixed parameters. Cells are arranged in three $24x24$ square layers. Only the first layer cells are reached by the input signal and only the last layer cells are used for output.

| **Parameter** | **Value** |
|---|---|
| Gene input fraction ($L$) | 0.5 |
| Reservoir genes ($N_r$) | 100 |
| Cells ($C$) | $1728(24 \cdot 24 \cdot 3)$ |
| Strains ($S$) | 10 |
| Input layers ($I$) | 1 |
| Output Layers ($O$) | 1 |
| ESM interaction distance ($\lambda$) | 15.0 |
| ESMs ($E$) | 5 |
| ESM threshold ($\theta$) | 11.5 |
| Delay ($d$) | 2 |
| ESM diffusion coefficient ($\beta$) | 5.0 |
| ESM basal secretion rate | 1.0 |
| ESM active secretion rate | 5.0 |
| Cell radius | 10.0 |
| Voxel side length | 20.0 |
| Warm-up time steps ($T_W$) | 100 |
| Total time steps ($T$) | 1000 |
| Function | *parity* |
| Window size ($w$) | 3 |

#### Supplementary figures

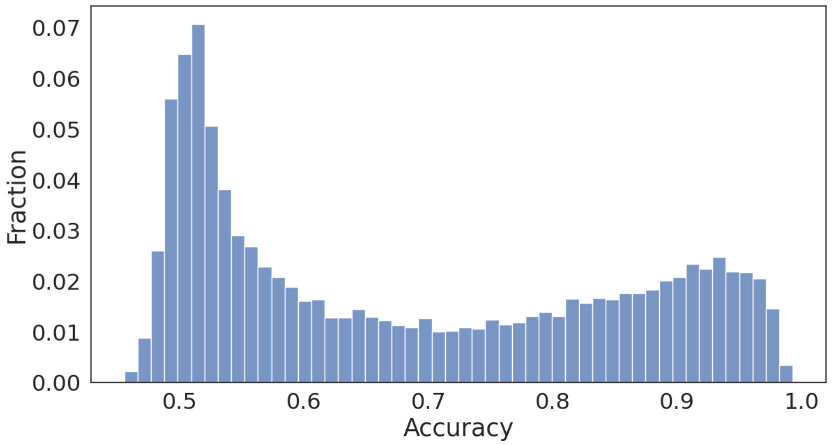


Figure S1: Simulation accuracy results for Sensitivity Analysis. Each of the 10,000 sets of parameters was simulated 5 times for a total of 50,000 simulations. The values for fixed parameters were chosen in a manner that would evenly distribute the accuracies between the 0.5 to 1.0 range.

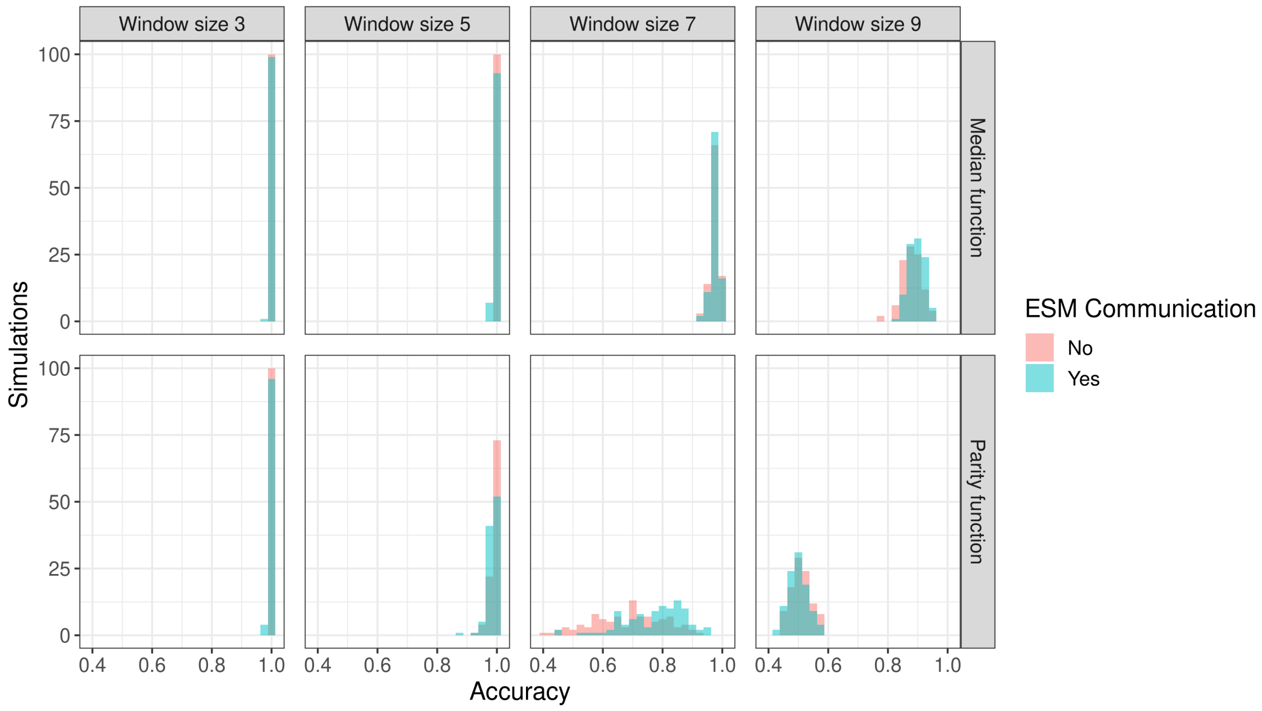


Figure S2: Comparison between two multicellular systems in which ESM communication is either allowed (bluegreen) or blocked (pink). Simulation parameters are the same as the Proof of Concept simulations (listed in Table S1) with the exception of ESM threshold $\theta$. Disabling ESM communication is achieved by setting the ESM threshold $\theta$ to 100000.0. Each histogram is the accuracy distribution of 100 simulations. As the histograms between the two systems mostly overlap, we can conclude that the ESM communication is not the major contributor to accuracy in the Proof of Concept scenario.

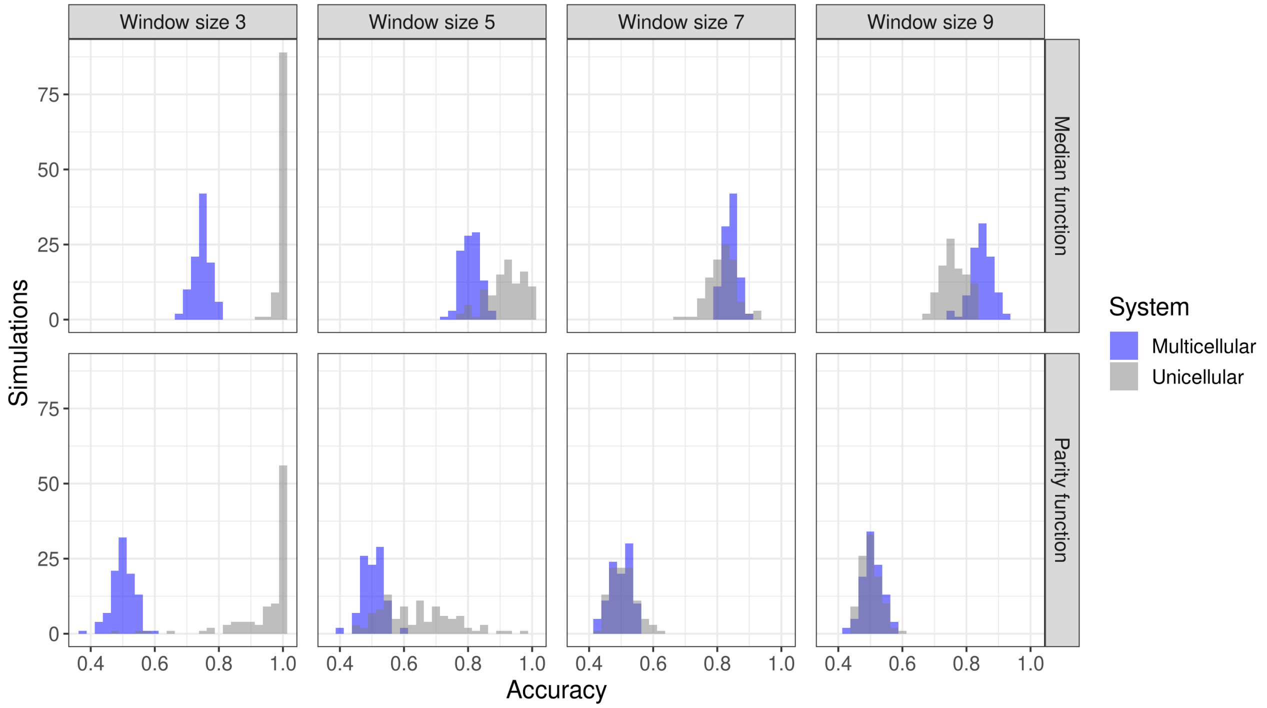


Figure S3: Comparison between a unicellular system and a multicellular system similar to the Proof of Concept analysis. Here, we have altered the multicellular system such that one half of the strains is capable of receiving the input signal and the other half is used for output. This analysis compares a unicellular system with a multicellular system in which ESM communication is necessary to carry information about the input signal without spatial heterogeneity. Each histogram is the accuracy distribution of 100 simulations. In contrast to the other scenarios, these multicellular systems improve in accuracy as the window size increases (visible in median function), indicating that memory is an influential component. Notably, unicellular systems perform better at shorter window sizes.

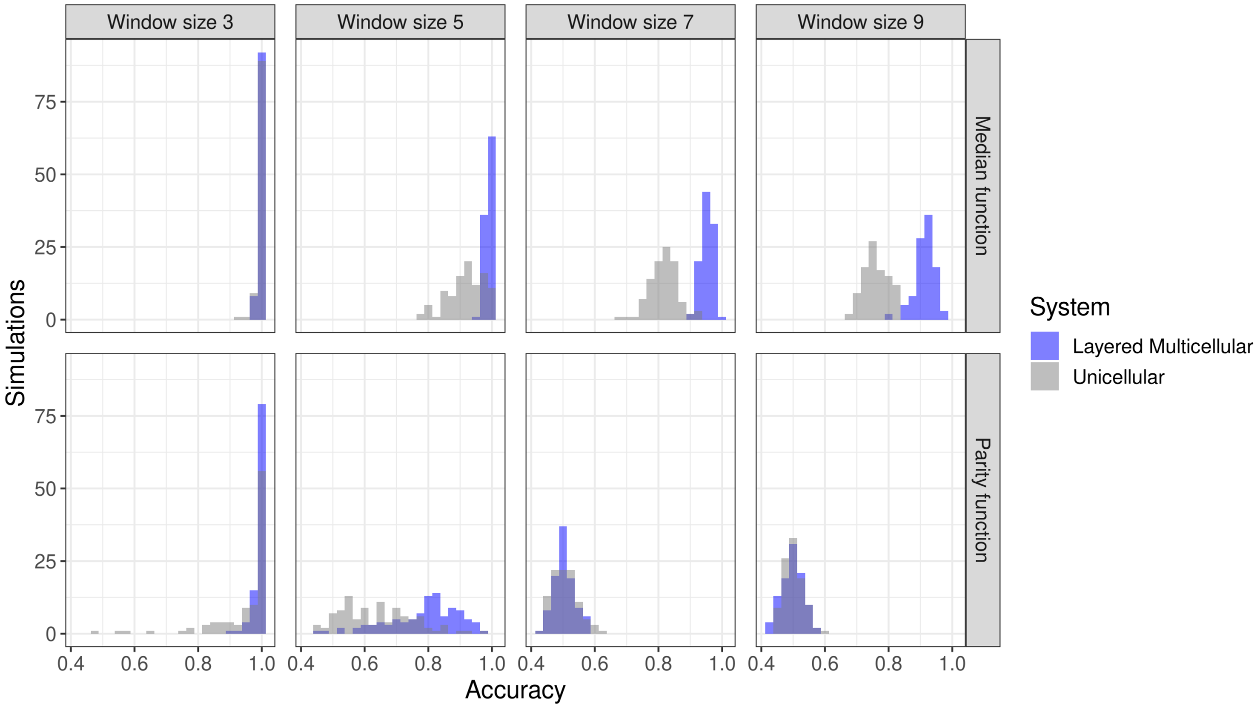


Figure S4: Comparison between a unicellular system and a Layered Community. In order to maximize accuracy and show the performance capabilities of the Layered community, the number of cells is 7500 and the number of ESMs 20. Each histogram is the accuracy distribution of 100 simulations. Layered Communities outperform unicellular systems, except when both perform equally poorly.

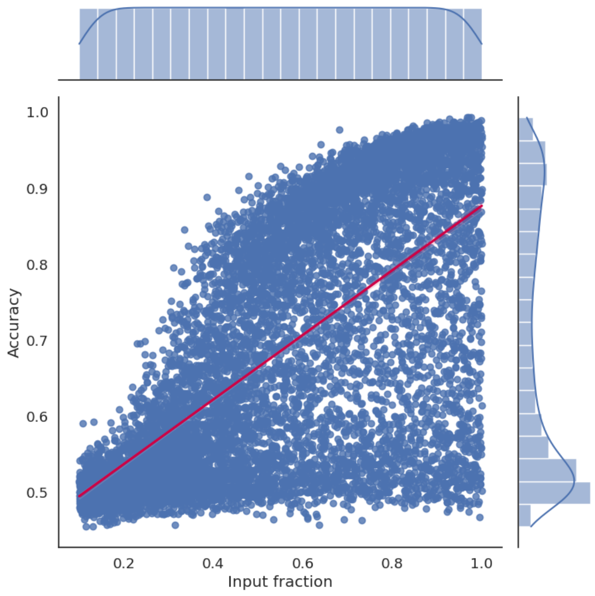


(a) Input fraction correlation

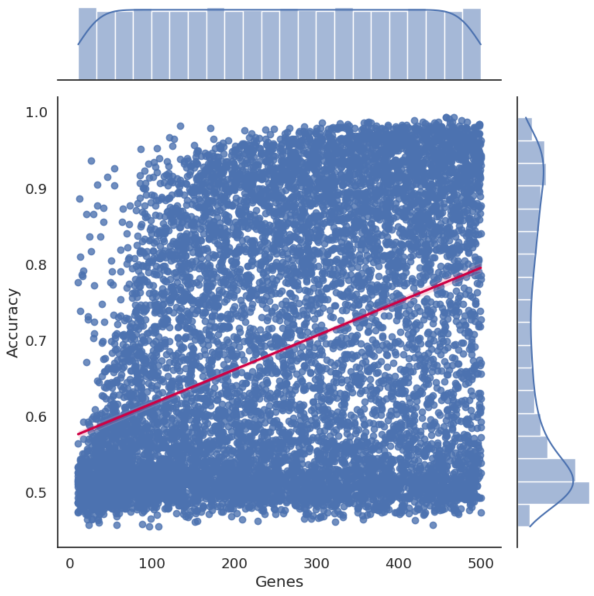


(b) Genes correlation

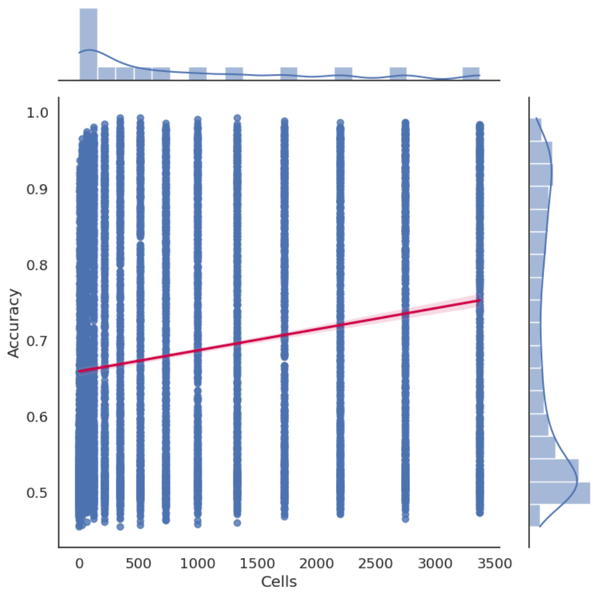


(c) Cells correlation

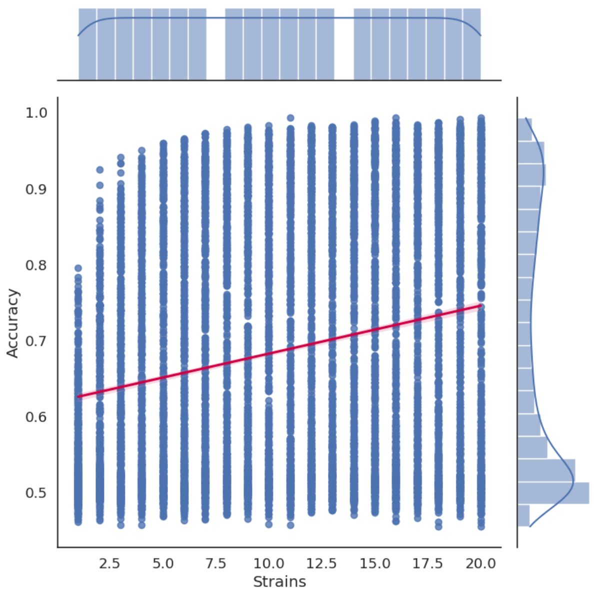


(d) Strains correlation

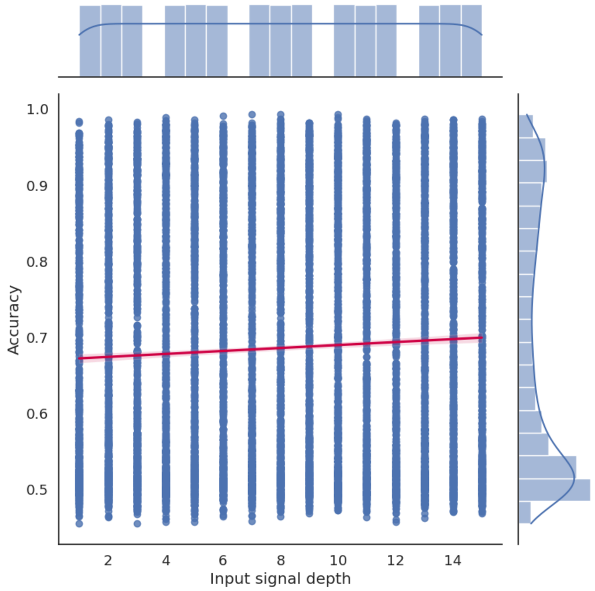


(e) Input layers correlation

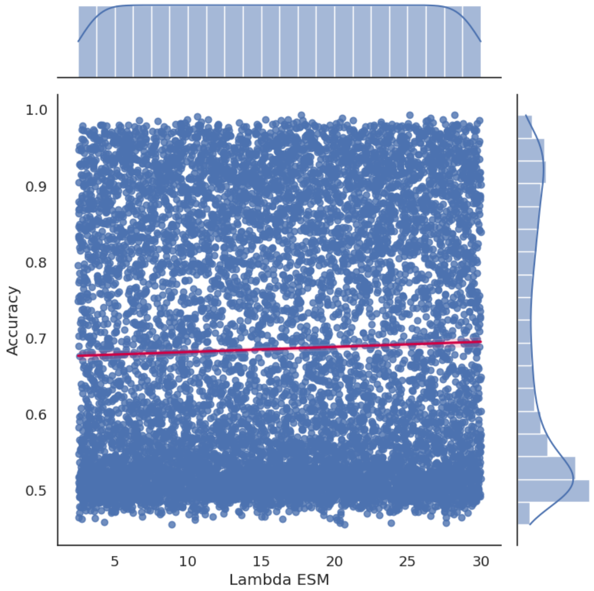


(f) ESM $\lambda$ correlation

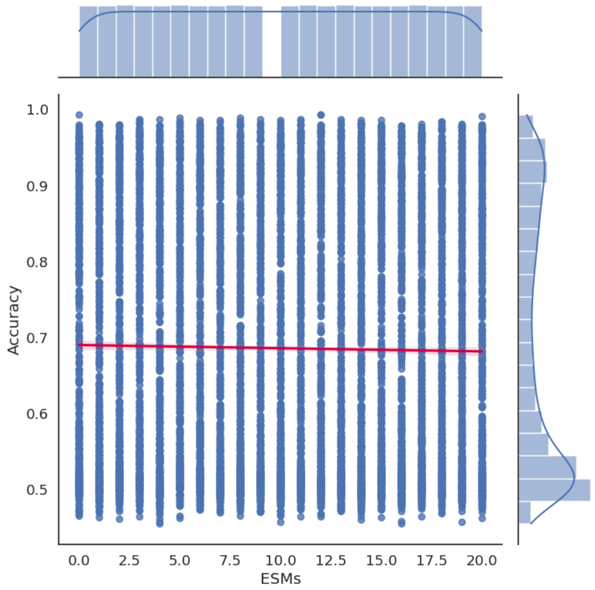


(g) ESMs correlation

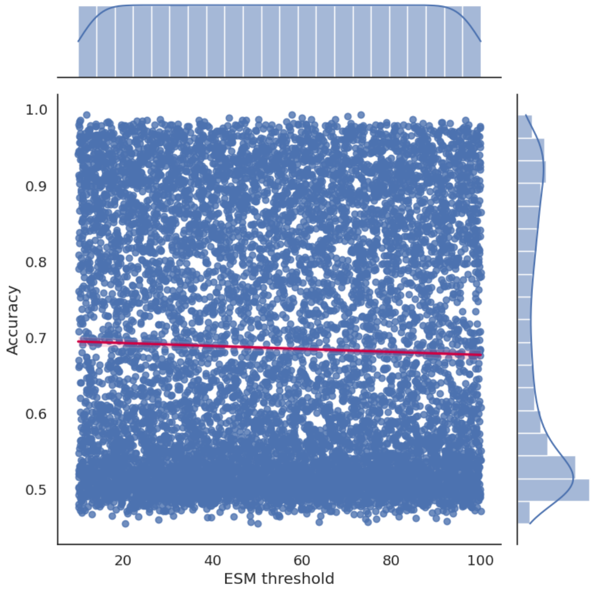


(h) ESM $\theta$ correlation

Figure S5: Sensitivity Analysis correlation plots for every tested parameter. The red line shows a linear regression fit. A total of 50,000 simulations were done.

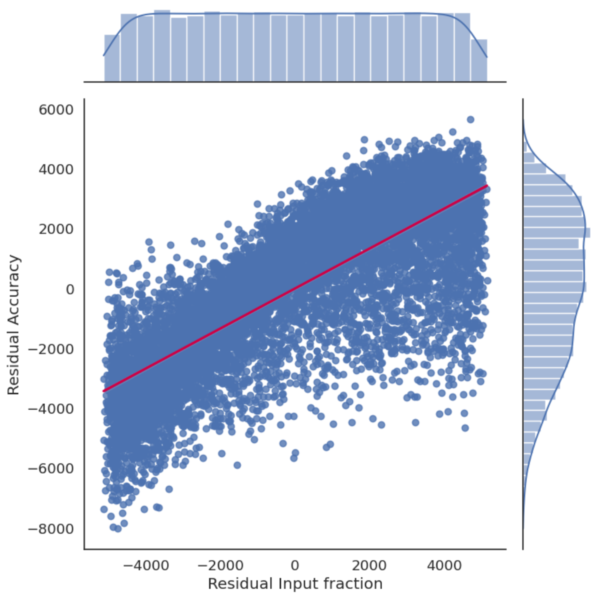


(a) Input fraction partial correlation

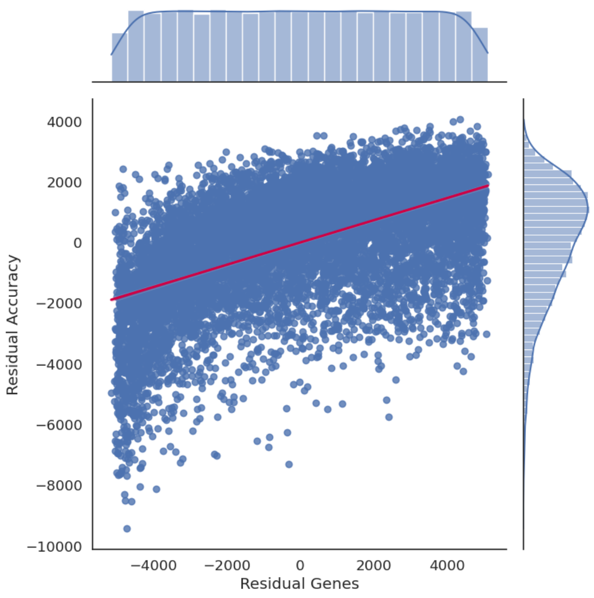


(b) Genes partial correlation

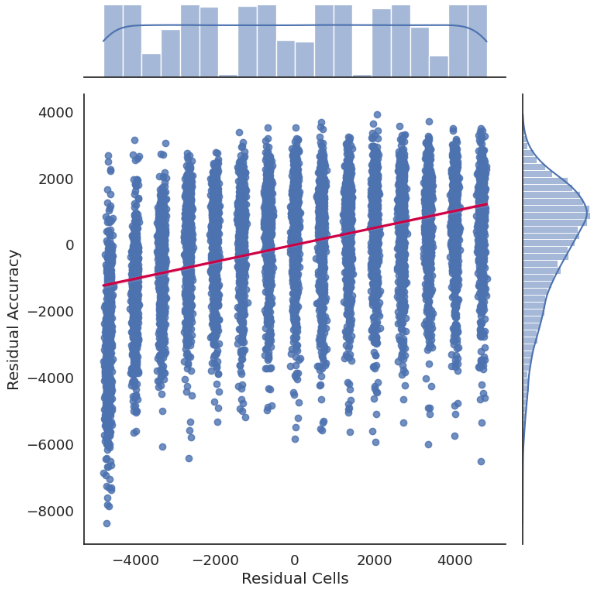


(c) Cells partial correlation

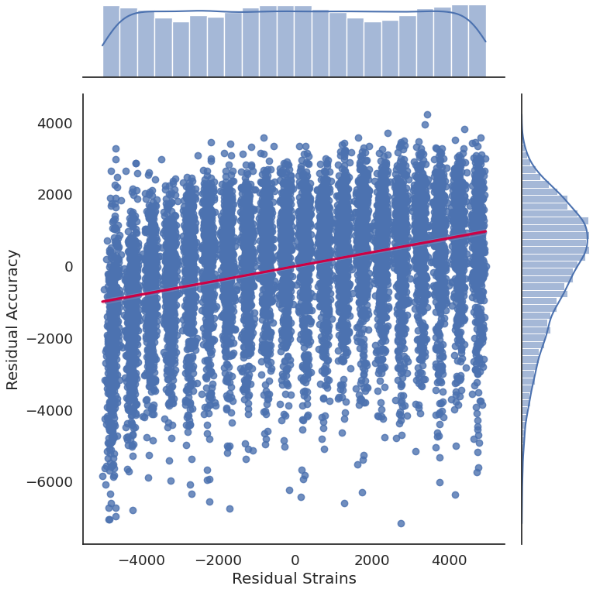


(d) Strains partial correlation

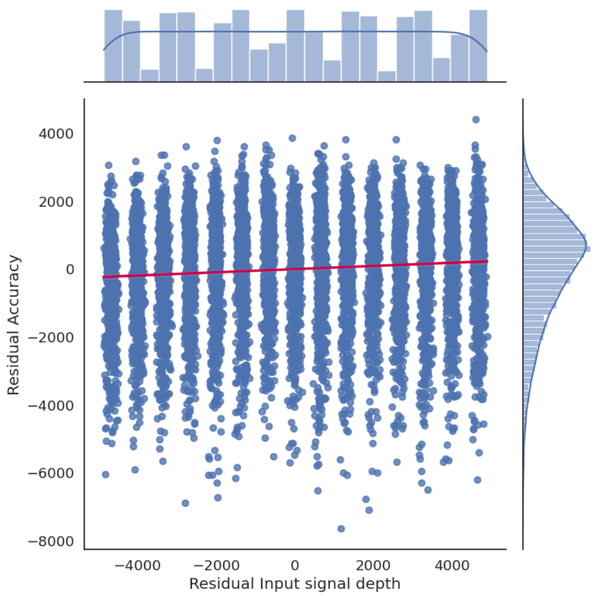


(e) Input layers partial correlation

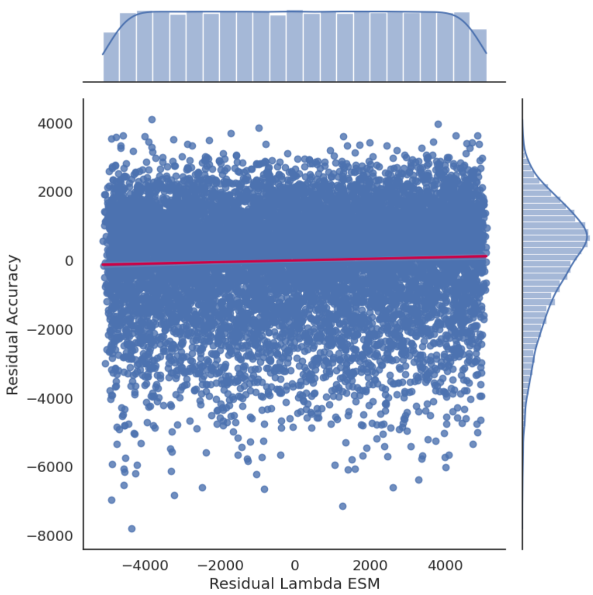


(f) ESM $\lambda$ partial correlation

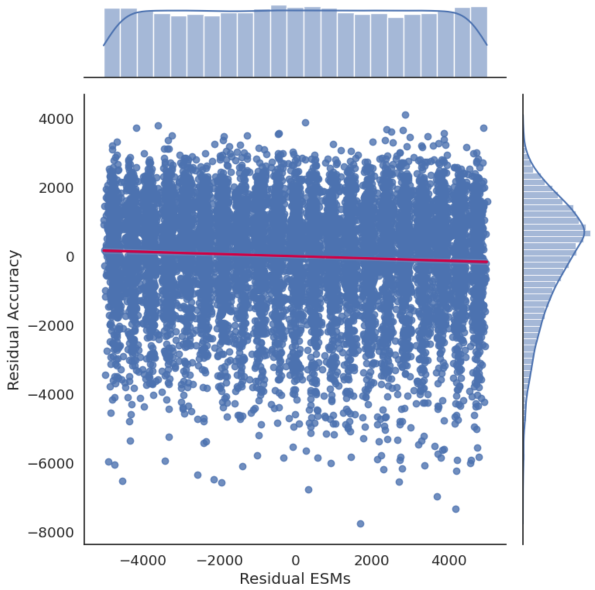


(g) ESMs partial correlation

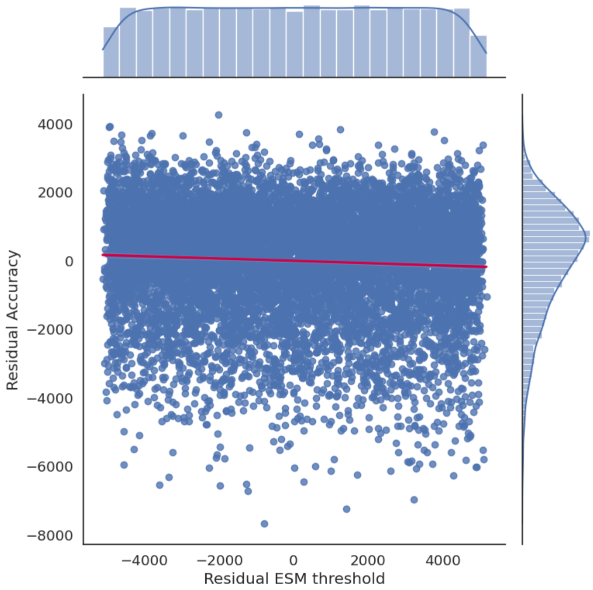


(h) ESM $\theta$ partial correlation

Figure S6: Sensitivity Analysis partial correlation plots for every tested parameter. The red line shows a linear regression fit. A total of 50,000 simulations were done.

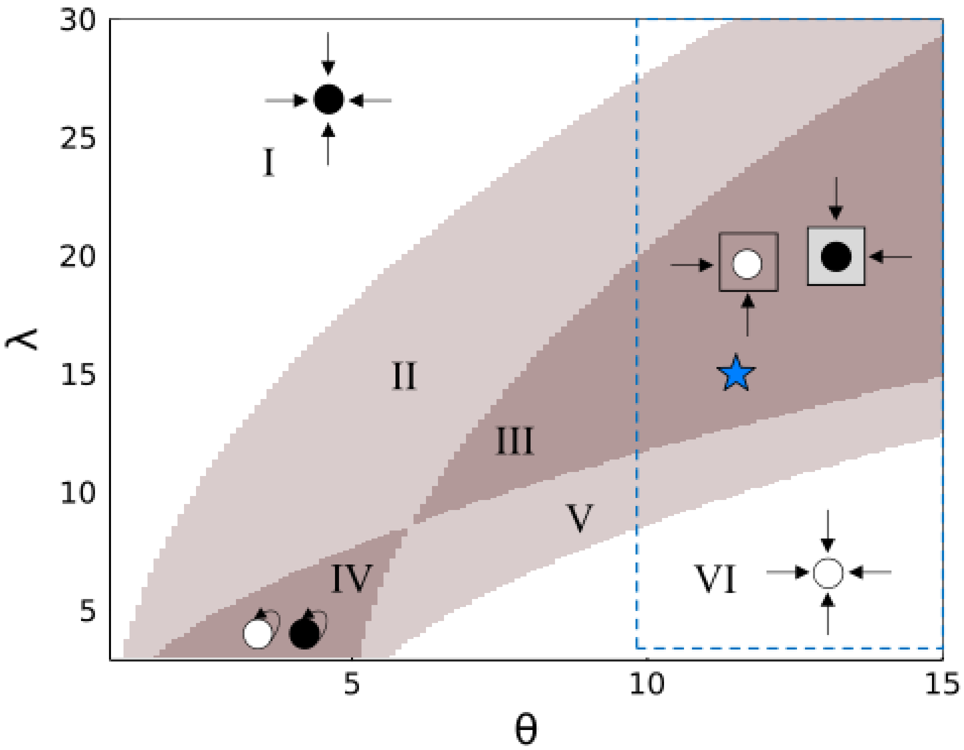


Figure S7: Communication regions within the $\lambda$, $\theta$ parameter space. In regions I, VI, and IV no communication occurs because the ESM receptor is always on, always off, or only responds to changes in self-generated signal, respectively. In regions II, III, and V communication between cells can occur; the receptor responds to either self- or neighbor-generated signal (II), only neighbor-generated signal (III), or the necessary combination of both self- and neighbor-generated signal (V). The blue star represents the $\lambda$, $\theta$ values chosen for the Proof of Concept and Layered Community analysis. Here, cells are communicating while limiting strong signal diffusion to the neighboring cells. The blue dotted box represents the range of parameter values used for Sensitivity Analysis. Within this range, all communication regions except IV are represented. Note that the range of $\theta$ extends beyond the plot ($\theta = 10 : 100$), but are not shown here to give a better view of the communication regions. This figure was originally published in Echlin(2019)(1) and has been superficially modified here.

### References

[1] Moriah Echlin. *A Complex Systems Approach to Understanding Cells as Systems and Agents*. PhD thesis, University of Washington, August 2019.
